# Ending AIDS: Progress and prospects for the control of HIV and TB in South Africa

**DOI:** 10.1101/061929

**Authors:** Brian G. Williams, Somya Gupta, Matthew Wollmers, Reuben Granich

## Abstract

We assess the prospects for ending AIDS in South Africa using a dynamical model to fit data on time trends in HIV prevalence and anti-retroviral treatment (ART) coverage for adults. We estimate current and project future trends in HIV incidence, prevalence and AIDS related deaths, in ART coverage and incidence, and in TB notification rates. We consider two scenarios: *constant effort* under which people continue to be started on treatment at the current rate and *expanded treatment and prevention* under which testing rates are increased, everyone is started on treatment as soon as they are found to be infected with HIV, and voluntary medical male circumcision, pre-exposure prophylaxis and condom distribution programmes are expanded.

As a result of the roll-out of ART the incidence of HIV has fallen from a peak of 2.3% *per annum* in 1996 to 0.65% in 2016, the AIDS related mortality from a peak of 1.4% *per annum* in 2006 to 0.37% *p.a*. in 2016 and both continue to fall at a *relative* rate of 17% *p.a*. Maintaining a policy of *constant effort* will lead to further declines in HIV incidence, AIDS related mortality and TB notification rates but will not end AIDS. Implementing a policy of *expanded treatment and prevention* in September 2016 should ensure that by 2020 new infections and deaths will be less than one per thousand adults and the UNAIDS Goal of Ending AIDS by 2030 will be reached. Scaling up voluntary medical male circumcision, pre-exposure prophylaxis and condom availability will avert some new infections but will save relatively few lives. Nevertheless, equity demands that people at very high risk of infection including commercial sex-workers, men-who-have-sex-with-men and young women should have access to the best available methods of prevention.

The current cost to the health services of managing HIV and TB among adults in South Africa is about US$2.1 Bn *p.a*. (0.6% of GDP *p.a*.) and this will rise to a peak of US$2.7 Bn *p.a*. in 2018 (0.8% GDP *p.a*.). As treatment is scaled up and prevention made available to those at high risk, the cost will fall to US$ 1.8 Bn *p.a*. in 2030 and US$ 1.0 Bn *p.a*. in 2050 as those that are living with HIV on ART, die of natural causes. The cost of testing people for HIV is never more than about 8% of the total cost and since testing is the *sine qua non* of treatment it will be essential to invest sufficient resources in testing. The cost of treating tuberculosis is never more than about 10% of the total and since this is the major cause of AIDS related illness and deaths, efforts should be made to optimise TB treatment.

Ending AIDS in the world will depend critically on what happens in South Africa which accounts for 20% of all people living with HIV. The increasing availability of ART has had a major impact on both HIV incidence and AIDS related mortality and universal access to ART is affordable. With the commitment to make treatment available to all those infected with HIV in September 2016, the South African government is well placed to eliminate HIV as a major threat to public health by 2020 and to end AIDS by 2030. Individuals at high risk of infection deserve access to the best available methods of protecting themselves and they will become increasingly important in the final stages of ending the epidemic.

## Introduction

The United Nations Joint Programme on HIV and AIDS (UNAIDS) has set a target to end AIDS in the world by 2030^1^ and mathematical models are needed to estimate current and project future trends in the epidemic of HIV, to determine the feasibility and cost of reaching the target, and to show what needs to be in place to ensure success. In particular it will be necessary to:

1. Agree on a definition of ‘the end of AIDS’;
2. Have access to reliable trend data on the prevalence of HIV and the coverage of anti-retroviral therapy (ART) and on tuberculosis (TB) notification rates, supplemented where possible by data on HIV incidence and mortality, rates of adherence to ART and of viral load suppression, and the prevalence of HIV in TB patients;
3. Agree on key aspects of the natural history of the epidemic which will determine the model structure to be used to fit trend data on the prevalence of HIV and the coverage of ART, and to estimate future trends in the incidence of HIV, AIDS related mortality and TB notification rates;
4. Agree on a suite of interventions for treatment and prevention as these will determine the prospects for reaching the UNAIDS target for 2030.
5. Agree on the costs of each intervention as well as the costs to society and individuals of HIV infection and AIDS deaths which will determine the overall costs, cost effectiveness and allocation of resources.

One in five people infected with HIV lives in South Africa and ending AIDS in the world depends on ending AIDS in South Africa. Managing the epidemic in South Africa represents an opportunity and a challenge: an opportunity to show what can be done and a challenge because of the scale of the problem in an uncertain health system. We use a dynamical model to fit trend data on the prevalence of HIV and the coverage of ART, estimate and project trends in HIV incidence and AIDS related mortality, and estimate current and future costs, under different combinations of treatment and prevention. We focus on adults but acknowledge that consideration must be given to eliminating vertical transmission and treating those that are infected vertically.^2–6^

## Model structure

The model, which has been described in detail elsewhere,^7^ is a standard susceptible-infected (SI) model with four stages of infection to reflect the Weibull survival distribution of adults who are not on ART^8^ and is not age-structured. Transmission falls as prevalence rises to reflect heterogeneity in the risk of infection.^7^ We allow for the possibility that changes in behaviour, in response to the rising epidemic, might have led to a fall in transmission but only if the data demand it. We assume that treatment is currently started only in those with low CD4-cell counts. The timing, rate of roll-out and asymptotic prevalence of ART are varied to fit the data.

As treatment coverage is expanded to include all those known to be infected with HIV we vary the timing, the rate of increase and the asymptotic testing rate so as to achieve 90% coverage of ART by 2020 and then continue to test people and start them on ART at the same rate. We allow for increased coverage of voluntary male circumcision (VMMC), pre-exposure prophylaxis (PrEP) and condom distribution.

The TB model is described in detail elsewhere.^7^ The model assumes that TB in HIV negative people is declining at a fixed rate and that the relative increase in TB among HIV positive people is driven entirely by the prevalence of HIV and the coverage of ART. The TB model has two variable parameters: the TB notification rate before the HIV epidemic started and the rate of increase of TB notification rates with HIV disease progression. We use notification rates rather than the incidence of TB because estimating the case-detection rate is fraught with difficulty and provided this does not change substantially over time the overall conclusions will not be affected.

## Data sources and assumptions

The trend data for HIV prevalence are taken from UNAIDS,^9^ the trend data for TB notification rates from the WHO Global Report on TB.^10^ Table 1 gives the assumptions concerning the relative rate at which people currently start ART in each clinical stage. Here we assume that, of those that test positive, everyone in Clinical Stage 4, half of those in Clinical Stage 3, one-quarter of those in Clinical Stage 2 and none of those in Clinical Stage 1, are started on treatment. Each of these proportions is multiplied by an overall rate at which people start treatment.

**Table 1.** The *relative* proportion of people starting ART in each of four clinical stages of infection, the proportion of those on ART that are virally suppressed and the relative transmission of those on ART as compared to those that are not on ART. ‘Current’ corresponds to the current treatment guidelines, ‘Expanded’ to the ‘treat-all’ guidelines to be implemented in September 2016.

| Clinical stages | Current | Expanded |
| --- | --- | --- |
| 1 | 0.00 | 1.00 |
| 2 | 0.25 | 1.00 |
| 3 | 0.50 | 1.00 |
| 4 | 1.00 | 1.00 |
| Suppressed | 0.90 | 0.97 |
| Transmission | 0.12 | 0.12 |

We consider two scenarios: *constant effort* under which South Africa continues to recruit people at the current rate and start treatment under the current guidelines; *expanded treatment and prevention* under which all those at risk are tested once every nine months, on average and anyone who is found to be infected with HIV is immediately started on ART.

Under the *constant effort scenario* we assume that 90% of people on ART are virally suppressed based on a study carried out in South Africa.^11^ Under the *expanded treatment and prevention* scenario we assume that 96.5% of people on ART are virally suppressed based on a study carried out in Botswana.^12^ We assume that the infectiousness of those that are not virally suppressed is 12% of the infectiousness of people that are not on ART (see Appendix 1).

East and Southern Africa refers to countries in each region as defined by UNAIDS. All rates and percentages are per adult aged 15 years or more.

## Fitting current trends

The model-fits to the trends in the prevalence of HIV and of ART are given in Figure 1A and fits to the trends in the TB notification rates in Figure 2E. The best-fit parameters are as follows. The HIV transmission parameter, which determines the initial rate of increase, is 0.706/year; the heterogeneity parameter which determines the peak prevalence is 0.242.^7^ The incidence of TB increases by 40% for each decline of 100 CD4^+^ T-cells per μL. Unlike the situation in most of the other countries of East and Southern Africa there is only slight evidence of a reduction in transmission over and above the natural history of the epidemic. We assume that the rate of starting ART increases logistically and the best fit parameters show that the overall asymptotic rate is 2.00/year, multiplied by the factors given in Table 1 for each clinical stage, reaching half the maximum value in 2011. For TB we assume that the rate of tuberculosis in HIV-negative people is falling by 1% *p.a*., that the incidence doubles during the acute stage of infection and then increases by a factor of 2.73 from one clinical stage to the next corresponding to an increase of about 49% for each decline in the CD4+ cell count of 100 cells/μL.

**Figure 1.**
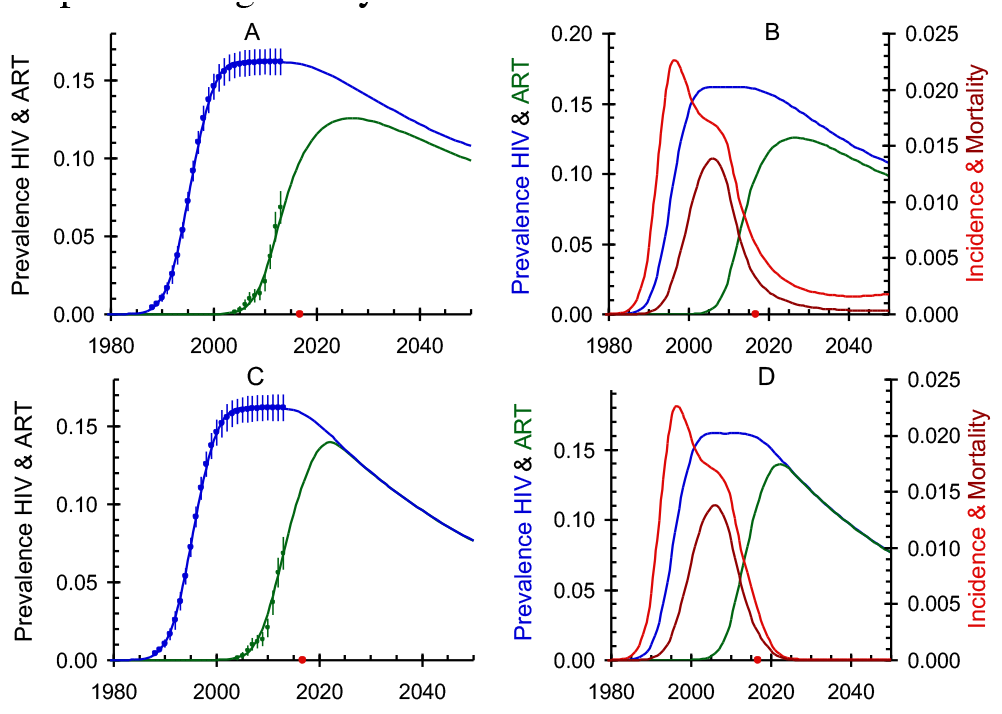
A and B: Constant effort; C and D: Expanded treatment and prevention. A and C. Blue: HIV prevalence; Green: ART prevalence in adults. B and D. Red: HIV incidence; Blue: HIV prevalence; Brown: AIDS related mortality; Green: ART prevalence.

**Figure 2.**
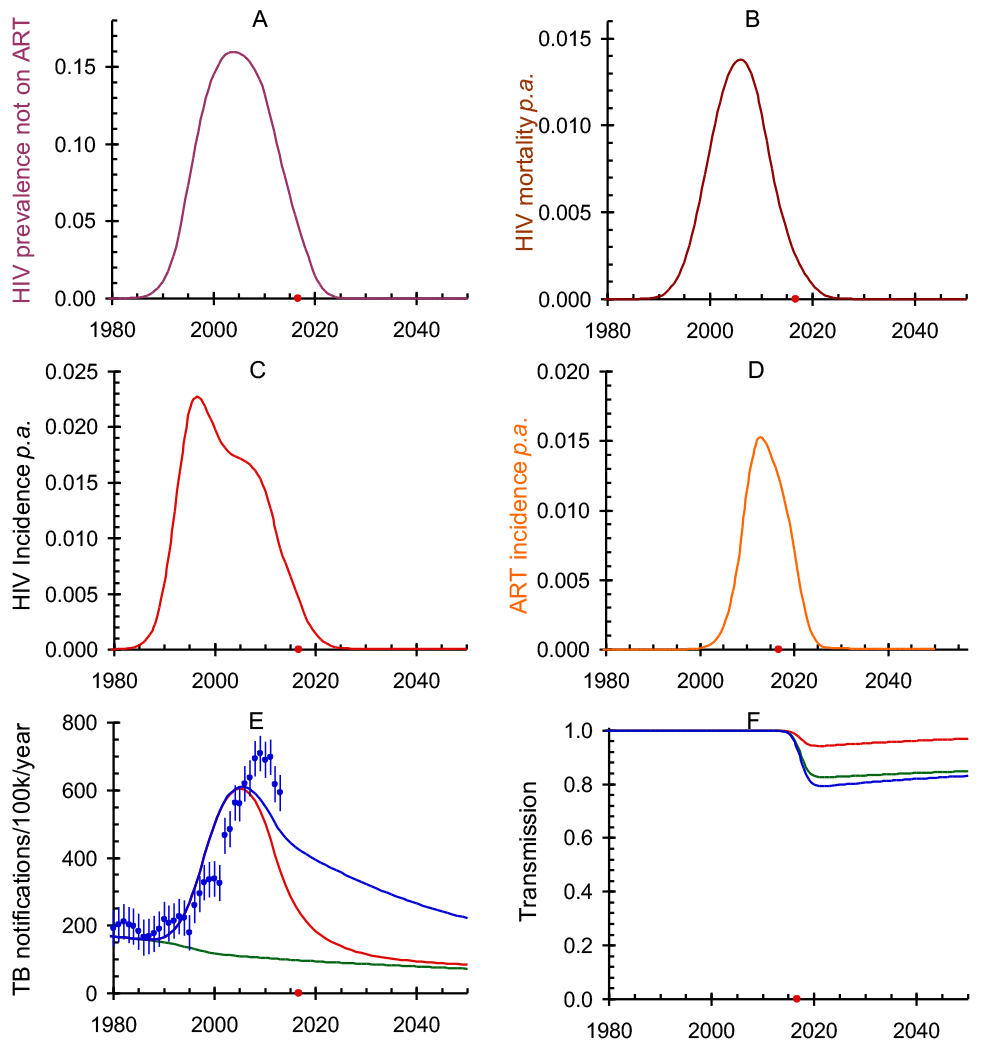
A: HIV prevalence not on ART; B: AIDS related mortality; C: HIV incidence; D: ART incidence; E: Tuberculosis notification rates. Green: HIV-negative; Red: Green plus HIV-positive not on ART; Blue: Red plus HIV-positive on ART. F: Reduction in transmission resulting from prevention. Red: VMMC Green: Red plus PreP; Blue: Green plus CD.

The model-fit to the prevalence of HIV and the coverage of ART is good (Figure 1A); the model-fit to the TB notification rates is less good (Figure 1B). The TB model has been shown to give good fits to trends in TB notification rates in most countries in Southern Africa^7^ as well as in East Africa (Williams *et al*. in preparation). Furthermore, the underlying trend data for the prevalence of HIV based on the annual ante-natal clinic surveys^13^ are reliable. One might wish to examine the reported TB notification rates in South Africa more carefully.

## Estimating current impact

Providing ART to those infected with HIV has had a major impact on the epidemic. An estimated 47% of all those living with HIV are now on ART and of these an estimated 92% had viral loads below 400/μL.^14^ As a result of the rollout of ART the incidence of HIV has fallen from a peak of 2.3% *per annum* in 1996 to 0.65% in 2016, the AIDS related mortality has fallen from a peak of 1.4% *per annum* in 2006 to 0.37% *p.a*. in 2016 and both continue to fall at a *relative* rate of 17% *p.a*. (Figure 1B). Although the prevalence of HIV has remained constant for the last ten years this is largely due to the increase in the number of people who are being kept alive on ART which balances the decline in the number of new infections. Continuing to provide ART under the current policy will not, however, bring about the End of AIDS by 2030 (Figure 1B).

## Expanding treatment

The South African government is committed to providing ART to all those found to be infected with HIV. Here we assume that by 2017 the conditions for starting ART are changed so that everyone who is infected with HIV is eligible for treatment (Table 1). We assume that voluntary medical male circumcision (VMMC), pre-exposure prophylaxis (PrEP) and condom distribution (CD) are rolled out as indicated in Table 2. For all three prevention interventions we assume that the expansion started in 2016 and full coverage is reached in 2020. For VMMC we set the current coverage to 46% and the target coverage to 80% with an HIV-incidence risk ratio for circumcised versus uncircumcised men of 0.4.^15^ For PrEP we set the current coverage to zero and the target coverage to 50% assuming that the proportion of women needing PreP is proportional to the overall prevalence of HIV. We set the HIV incidence risk-ratio for women on PrEP to 0.4.^15^ For CD we set the current coverage to zero and the target coverage to 50% assuming that the proportion of men needing condoms is proportional to the overall prevalence of HIV. We set the HIV incidence risk-ratio if men use condoms to 0.46 which allows for low adherence rates.^15^

**Table 2.** For each intervention the table gives: the year in which treatment expansion starts and when it reaches full coverage, the current coverage, the target coverage and the risk ratio for those receiving each intervention. VMMC: voluntary medical male circumcision; PrEP: pre-exposure prophylaxis; CD: condom distribution.

|  | VMMC | PrEP | CD |
| --- | --- | --- | --- |
| Start year | 2015 | 2015 | 2015 |
| End year | 2020 | 2020 | 2020 |
| Current coverage | 0.46 | 0.00 | 0.00 |
| Target coverage | 0.80 | 0.50 | 0.50 |
| Risk ratio<sup>15</sup> | 0.40 | 0.20 | 0.46 |

With these assumptions we project the HIV prevalence (Figure 1A), HIV prevalence of those not on ART (Figure 2A), AIDS related mortality (Figure 1B and Figure 2B), HIV incidence (Figure 2C), ART incidence, that is to say the proportion of people who start ART each year, (Figure 2D) TB notification rates (Figure 2E), and the cumulative reduction in transmission attributable to each of the prevention interventions (Figure 2F). The UNAIDS estimates of incidence, based on the same data for HIV-prevalence and ART-coverage are significantly different from ours, as discussed in Appendix 2.

Comparing Figure 1A and 1B with Figure 1C and 1D shows the additional impact of expanded treatment and prevention. By 2020 almost all those infected with HIV will be on treatment and there will be a small but significant additional decline in the TB notification rates.

## Epidemiological impact

Starting in September 2016 everyone in South Africa who tests positive for HIV will be eligible for ART. Here we assume that testing is scaled up so that by 2020 those at risk of HIV are tested, on average, once every nine months and the proportion of those on treatment will reach 96.5% as is already the case in Botswana.^12^ By 2030 AIDS related mortality and the incidence of HIV and in adults will fall to less than one new case and one death per thousand adults (Figure 2B and C). While the number of people on ART will remain high, decreasing slowly as those on ART die of natural causes (Figure 1D), the rate at which new people start ART will be very low after 2030 (Figure 2D).

The increased coverage of ART will bring down the TB notification rate but because ART only reduces the increased risk of TB by 61% (54%–68%)^16^ it will remain substantially higher than the pre-ART rate unless ways can be found to reduce TB in HIV-negative people more quickly (Figure 1D).

## Projecting costs

In order to project the future costs of controlling HIV we use cost estimates, given in Table 3, from the South African HIV and TB Investment Case^17^. ‘Testing’ includes the cost of delivering the test; ‘ART’ includes the cost of delivering treatment; ‘PrEP’ includes the cost of delivering the drugs; ‘Condoms’ includes the cost of condoms assuming that a man uses an average of 100 condoms each year.

**Table 3.** Costs in 2016 US$ for the provision of health care for those off and on ART for each of four clinical stages, delivery of tests, ART, VMMC, PrEP, condom distribution, deaths of young adults and TB treatment, and the Gross Domestic Product.^17^ We assume a discount rate of 3% *p.a*.

| Health care | US\$ | Treatment & prevention | US\$ |
| --- | --- | --- | --- |
| Cost of health care per person not on ART by clinical stage |  |  |  |
| 1 | 25 | Testing per test | 32 |
| 2 | 51 | ART per person *p.a.* | 274 |
| 3 | 63 | VMMC per circumcision | 101 |
| 4 | 108 | PrEP per person *p.a.* | 84 |
| Cost of health care per person on ART by clinical stage |  |  |  |
| 1 | 48 | Condoms per person *p.a.* | 5.67 |
| 2 | 54 | Deaths per person | 1.1k |
| 3 | 106 | TB treatment per case | 780 |
| 4 | 132 | GDP *per capita* | 7.6k |

For ART we use estimates of the cost of drugs plus delivery, care and support, for health care on and off ART as given in the South African Investment Case Report.^17^ For tuberculosis the cost per patient treated is the total number of notified cases divided by the total cost of the TB programme in 2014.^10,18^

Using the costs in Table 3 we estimate the programme costs over time, allowing for a discount rate of 3% p.a. (Figure 3). The cost of VMMC peaks early on as the backlog of uncircumcised men is made up, after that one only needs to circumcise men at the rate at which boys reach adulthood. The cost of PrEP and condoms both peak in 2020 but then decline as the prevalence falls and the number of women needing PrEP and men needing condoms fall with it.

**Figure 3.**
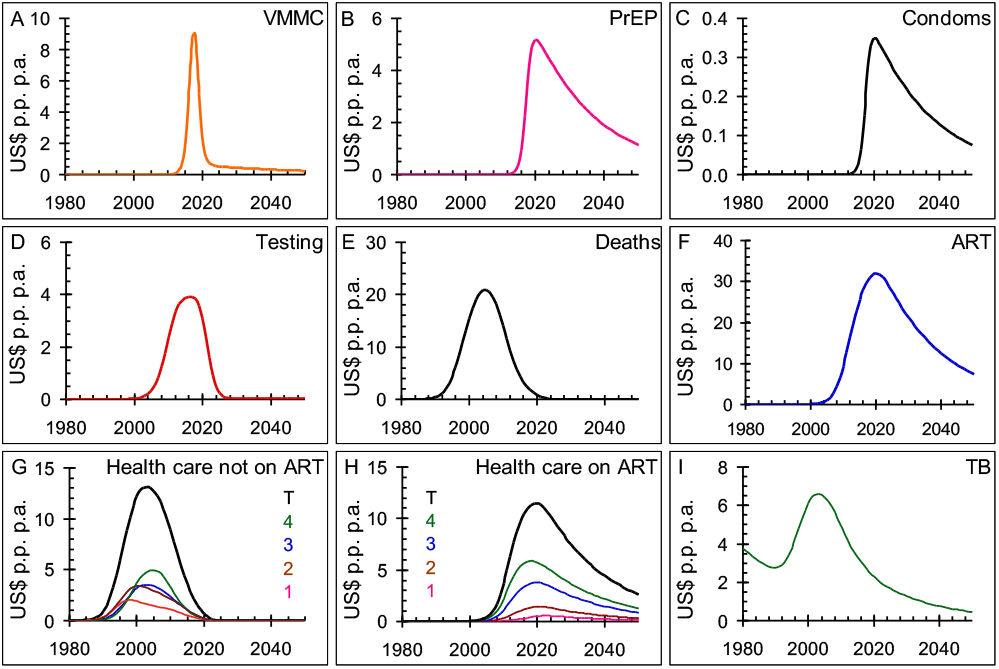
Cost of interventions in 2016 US$ per adult per annum. A: Voluntary medical male circumcision; B: Pre-exposure prophylaxis; C: Condom distribution; D: Testing for HIV; E: Deaths of young adults; F: Provision of ART; G: Health care for those not on ART by clinical stage and total; H; Health care for those on ART by clinical stage and total; I: Treatment of tuberculosis.

With random testing one would need to test 1/*P* people, where *P* is the prevalence of HIV, to find one person infected with HIV. As the prevalence falls testing will become more focused and may rely on contact tracing. We therefore let *N*(*i*), the number of people that are tested at time *t* for each person found to be infected with HIV, be

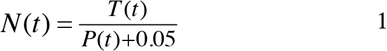

where *T*(*t*) is the proportion of people that start treatment and *P*(*t*) is the prevalence of HIV at time *t*. It would never be necessary to test more than 20 people to find one infected person.

Figure 4 and Figure 5 give the cost of managing HIV and treating TB, per adult and in total, over time where we have included the different interventions in the order indicated. Assessing the cost to society of letting young, working age adults die is hard. Here we set the cost to the health services of each AIDS death to 15% of the *per capita* gross domestic product (GDP) as discussed in Appendix 3. In 2005 the cost of deaths due to AIDS peaked at about US$700M *p.a*, after that it starts to fall as ART is rolled out and the major cost becomes that of providing ART which is currently running at about US$ 1.1Bn *p.a*. Under the scenario outlined here the total cost will peak at about US$2.7Bn*p.a*. in 2018.

**Figure 4.**
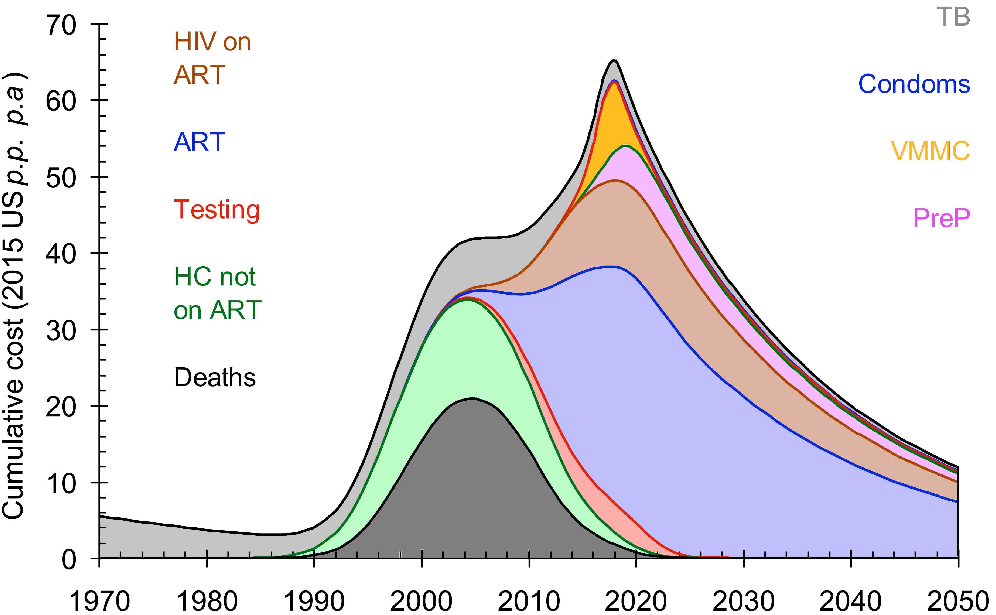
Cumulative costs of the different interventions in US$ per adult person per annum. Bottom to Top. Black: Deaths of young adults; Green: Health care for those not on ART; Red: Testing for HIV; Blue: Provision of ART; Brown: Health care for those on ART; Pink: Pre-exposure prophylaxis; Orange: Voluntary medical male circumcision; Blue: Condom distribution (hidden between VMMC and TB); Grey: Treatment of tuberculosis.

**Figure 5.**
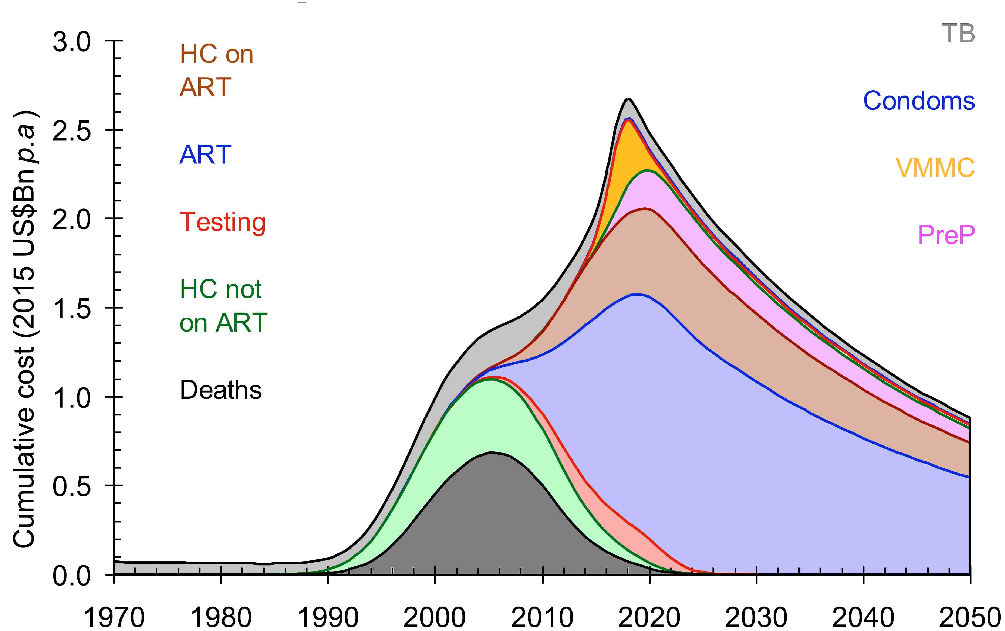
Cumulative total costs of the different interventions. Bottom to Top: Bottom to Top. Black: Deaths of young adults; Green: Health care for those not on ART; Red: Testing for HIV; Blue: Provision of ART; Brown: Health care for those on ART; Pink: Pre-exposure prophylaxis; Orange: Voluntary medical male circumcision; Blue: Condom distribution (hidden between VMMC and TB); Grey: Treatment of tuberculosis.

The cost of providing health care to those not on ART is substantial but only until 2020. The cost of testing is never more than about 10% of the total costs and since testing is the *sine qua non* of treatment these costs must be met. The cost of providing health care to those on ART is typically about 15% of the cost of ART and these costs could be combined. The cost of PrEP is relatively small; the cost of VMMC is significant but only for about five to ten years as the backlog of uncircumcised men is made up. The cost of condom promotion is very small and the cost of TB treatment is never more than about 5% of the total costs once the epidemic of HIV has become established.

The total cost peaks at about US$70 per adult *p.a*. in 2018 but falls rapidly thereafter to US$33 per adult *p.a*. in 2030 and US$12 per adult *p.a*. in 2050. These numbers are to be compared with the GDP for South Africa which is currently about US$7.6k per person *p.a*. or roughly double that if expressed as the GDP per adult *p.a*. The total cost is therefore never more than 0.5% of GDP.

In absolute terms the total cost will peak at about US$2.7Bn *p.a*. in 2020 and will fall steadily after that. In 2013 total spending on HIV and TB in South Africa was ZAR22Bn^17^ or US$2.3Bn taking the exchange rate, averaged over the year, of 9.5 ZAR/US$. Our estimate for 2013 is US$1.8Bn. If we assume that overhead costs and costs associated with HIV in infants and children are about 25% of the total costs, our estimates and the total expenditure are in good agreement.

It is important to compare the cost and benefits of *constant effort*, in which South Africa continues to provide ART under the previous guidelines and at the present rate, with the cost and benefits of *expanded treatment and prevention* as presented here. In Figure 6 we plot the number of new infections, AIDS-related deaths and the costs of the various combinations of interventions between 2016 and 2050. If South Africa were to continue with the pre-2016 guidelines and rate of scale up of treatment one would expect to have 10.4 million new HIV infections and 6.0 million AIDS related deaths at a cost of US$44 Bn. Implementing the new treatment guidelines will avert 3.6 million new infections, save 1.3 million lives and save US$1.3 Bn. If we add VMMC this will avert a further 113k new infections, 2.6k lives but will cost an additional US$1.3 Bn at US$12k per infection averted and US$104 per life saved. If we add PrEP this will avert a further 48k new infections, 12k lives at US$82k per infection averted and US$1.7M per life saved. If we add condom distribution this will avert a further 31k new infections and 800 lives at US$5.2k per infection averted and US$186k per life saved.

**Figure 6.**
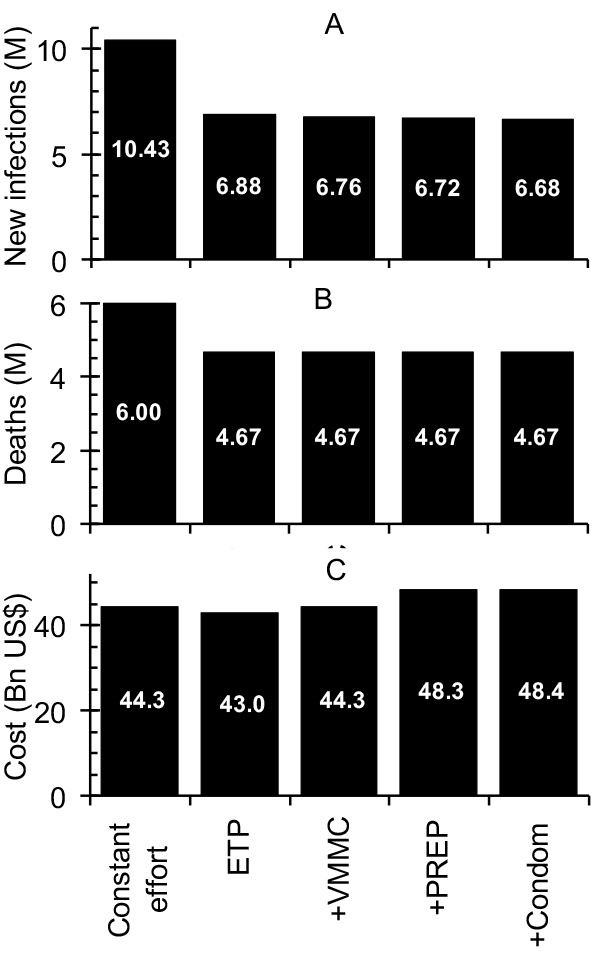
The cumulative number of A: new HIV infections, B: deaths, and C: the total cost of the HIV and TB programmes from 2016 to 2050 assuming (left to right) that South Africa continues to roll-out ART at the present rate (*constant effort*), or with *expanded treatment and prevention* (ETP) and, considered in succession: roll-out of voluntary medical male circumcision, PrEP, and condom distribution.

## Discussion

South Africa has the greatest number of people living with HIV and the greatest number on ART of any country in the world. As a result of the successful roll-out of ART the incidence of HIV among adults has fallen, from 2.3% *p.a*. in 1996 to 0.65% *p.a*. in 2016, a decline of 72%, and AIDS related mortality from a peak of 1.4% in 2006 to 0.37% in 2016 a decline of 68%. If South Africa expands treatment eligibility to all those found to be infected with HIV, invests sufficient funds in testing, and provides appropriate prevention interventions for those at high risk of infection, this will reduce HIV incidence and AIDS related mortality to less than one per thousand people by 2020 and reach the UNAIDS target to end AIDS by 2030.

HIV and TB currently cost South Africa about US$2.1 Bn (0.6% of GDP) and this will rise to a peak of US$2.7 Bn in 2018 (0.8% GDP). As treatment is scaled up and prevention made available to those at high risk it will fall to US$ 1.8 in 2030 and US$ 1.0 Bn in 2050 as those living with HIV on ART die naturally.

The cost of testing is never more than about 8% of the total cost and since testing is the *sine qua non* of treatment it will be essential to invest sufficient resources in testing. The cost of treating tuberculosis is never more than about 10% of the total and falling and since this is the major cause of AIDS related illness and deaths efforts should be made to optimise TB treatment.

If the treatment targets are met this will effectively eliminate AIDS related deaths so that expanded prevention will not save significantly more lives. Between 2016 and 2050 expanding treatment will avert 3.6 million infections, save 1.3 million lives and save US$1.3 Bn. If all the prevention methods are added to expanded treatment they will not save significantly more lives. However, VMMC will avert 113k new infections at a cost of US$12k per infection averted, adding PrEP will avert a further 48k new infections at a cost of US$82k per infection averted and scaling up the distribution of condoms will avert a further 31k new infections at a cost of US$5.2k per infection averted.

Equity demands that those at very high risk of infection have access to the best possible methods of protecting themselves. For people among whom the annual incidence is greater than 10% *p.a*., say, then if we assume that PrEP reduces the risk of infection by about 50% at a cost of the order of US$100 *p.a*. the cost of averting one infection in one person would be about US$2k. If we assume that VMMC reduces the risk of infection by 60% but the risk of infection over ten years is, say, 80%, then the cost of averting one infection in one person would be about US$250 per circumcision.

All three prevention interventions only contribute a small part of the overall cost. If PrEP is targeted at those at high risk it will also be cost effective. While the additional impact on HIV of VMMC will be relatively small^19^ in the context of *expanded treatment* it has many benefits in addition to reducing men’s risk of acquiring HIV: it protects men against a range of sexually transmitted infections^20,21^ and women from acquiring human papilloma virus which can lead to cervical cancer.^22–24^ It is, of course true, that if the various methods of prevention were rolled out without any increase in treatment coverage, the impact of prevention would be correspondingly greater.^28,29^

Finally, we note that in a recent paper on *The anticipated clinical and economic effects of 90–90–90 in South Africa*, Walensky *et al*.^25^ arrive at somewhat different results from ours (Appendix 2). Comparing the predictions of our model over the next ten years with the Walensky model^25^ their *current pace strategy* model predicts about twice as many new infections and six times as many deaths as our *constant effort model* while their *UNAIDS strategy* predicts about predicts about six times as many new infections and twelve times as many deaths as our expanded treatment and prevention strategy. Furthermore, the estimated total cost, over the next ten years, in the Walensky model^25^ is about twice that of our model (Appendix 5) and these differences need to be resolved.

Similar models for Mozambique^26^ and South Africa^27^ have predicted different outcomes depending on the parameterization. In these studies the critical parameter is the proportion of people on ART that are not virally suppressed. The study on Mozambique^26^ assumes that under their *accelerated scale-up* only 85% of those with a CD4+ cell count below 350/*μ*L, or about 50% of all those infected with HIV, are on ART and that ART reduces infectivity of HIV patients by only 80%. The study on South Africa^27^ assumes that only 77% (55%–94%) of those on ART are virally suppressed. It goes without saying that if ART coverage is low and that if many of those on ART are not virally suppressed it will be difficult to end AIDS.

## Conclusions

The effectiveness of modern anti-retroviral drugs is such that with a new commitment to getting as many people as possible onto ART while ensuring high levels of compliance and effective viral load suppression it will be possible to end AIDS by 2030 in South Africa, the country that accounts for about 20% of all those living with HIV.

Expanding access to testing and treatment is both feasible and cost effective and will result in significant reductions in transmission, illness and deaths while greatly reducing the burden on the health services. In the context of expanded treatment, prevention will have a relatively small additional impact. However, as transmission in the whole population declines there will still be significant numbers of infections in small and possibly hard to reach populations. For these people good prevention interventions may be critical in bringing about the ‘End of AIDS’. The ‘End of AIDS’ does not of course mean the end of HIV and South Africa will have a significant number of people alive and infected with HIV for the next fifty years, or until a cure is found, and these people will need care and support.

Program performance matters and it will require considerable focus, an efficient service delivery model to ensure early diagnosis and sustained treatment, and good surveillance to monitor progress and identify and deal with any weaknesses. The key markers of success will be the proportion of all those infected with HIV that are on ART and the viraemia in those on ART. The most important weakness in our ability to assess progress and project the future of the epidemic remains the lack of good surveillance and patient monitoring systems in many of the worst affected countries. Models are only as good as the data that informs them and greater efforts must be made to collect, assemble, synthesize and analyze much better data on surveillance and patient monitoring.

## Appendix 1: Proportion suppressed and transmission in those not virally suppressed

We need to decide on the proportion of people failing ART, *ϕ*, and the mean viral load among those on ART and failing or not failing treatment which could be set to, say, *ν_f_* = 1000/*μ*L and *ν_a_* 100/*μ*L. We then calculate the mortality and transmission among those failing treatment. We use the following relationships between viral load and mortality and transmission based on data from the distribution of viral loads in a study in Orange Farm and the relationship between viral load and survival^30^ to determine the relationship between the survival in years, *σ*, and the viral load per *μ*L, *ν*.

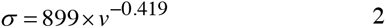

Including a background mortality of 2% *p.a*., unrelated to HIV, we set the mortality rate for those infected with HIV to be

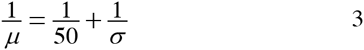

We use data from the relationship between viral load and infectiousness to determine the following relationship between infectiousness and viral load^31,32^

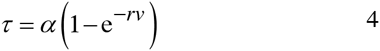

with *α* = 0.0858 and *r* = −1.583×10^−4^. With these values the relationship betwee viral load, life-expectancy and transmission is as given in Figure 7.

**Figure 7.**
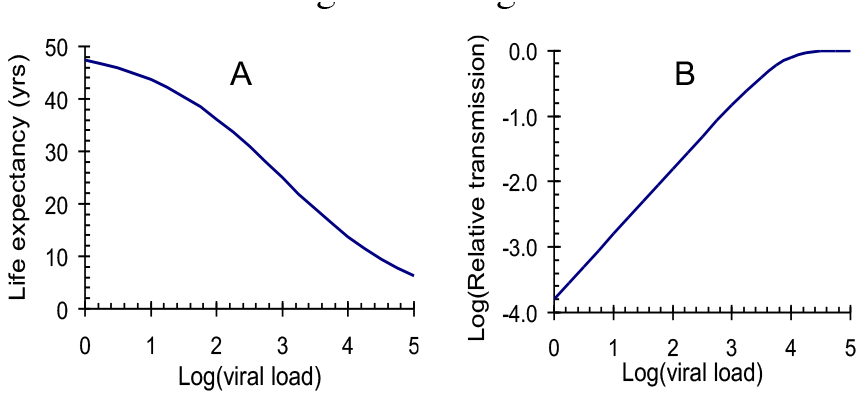
The life expectancy (A) and relative transmission rate (B) as a function of viral load.

We need to know the proportion of people who fail to suppress their viraemia, their life expectancy and their relative infectiousness. A recent study from Botswana^12^ showed that, of those on ART, 93.1% (92.1%–94.0%) had a viral load below 40/μL and 96.5% (96.0%–97.0%) had a viral load below 400/μL. If we assume that the average viral load of those whose vireamia is less than 40/μL was 20/μL then their infectiousness is 0.003 times that in those not on ART. If we assume that the average viral load on those below 400/μL was 200/μL then their infectiousness is 0.031.times that in those not on ART. The average infectiousness of those on ART is then 0.004 times that in those on ART. If we assume that the average viral load of those whose viraemia wass above 400/μL was 800/μL then their infectiousness is 0.12 times that in those not on ART. We therefore assume that:

1. 92% of those on ART are virally suppressed;
2. In those that are virally suppressed transmission is reduced by a factor of 0.004 and life expectancy is 41 years
3. In those that are not virally suppressed transmission is reduced by a factor of 0.12 and life expecancy is 26 years.

We then have *μ_n_, μ_p_, μ_a_, μ_f_* and *τ_a_, τ_f_* for the mortality and transmission among those that are HIV-negative, HIV-positive, on ART and failing ART. Given *v_a_* and *v_f_* we can calculate *μ_n_, μ_p_, μ_a_* and *μ_f_* and we set *μ_n_* to 0.02 (for 50 years life expectancy), *μ_p_* to 0.1 (for ten years life-expectancy). We then calculate the mean transmission for those on ART as (1−*ϕ*)*μ_a_* + *ϕμ_f_*. and the mean transmissibility for those on ART as (1−*ϕ*)*μ_a_* + *ϕμ_f_*

With these assumptions the mean transmission for those on ART, compared to those not on ART, is

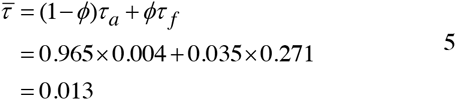

The mean annual mortality for those on ART is

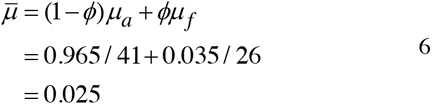

so that the life expectancy for those on ART is 40 years.

## Appendix 2: Comparison with UNAIDS model

We use the estimated prevalence of HIV over time as provided by UNAIDS which provides the input to the Spectrum/EPP Model. It should be noted that the estimated incidence derived from the trend in prevalence in the Spectrum/EPP Model differs from the one presented here as shown in Figure 8. Until 1995 the estimates are close but the UNAIDS estimate peaks at a higher incidence, falls rapidly between 1995 and 2009 and then levels off.

**Figure 8.**
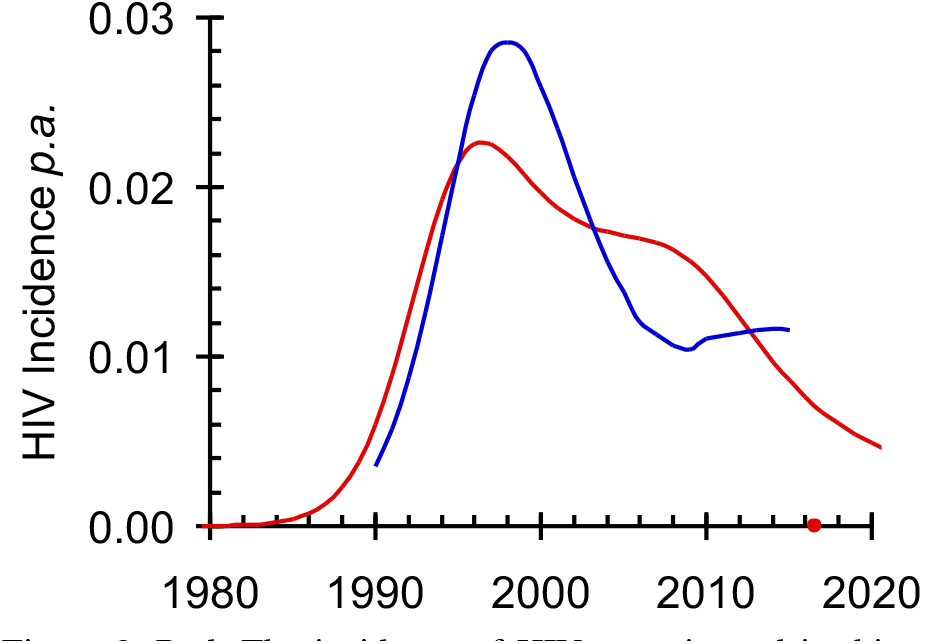
Red: The incidence of HIV as estimated in this paper; Blue: The incidence of HIV as estimated by UNAIDS.^9^

A partial explanation for the differences in the estimates in Figure 8 may be found in a study of HIV in Mozambique using the Spectrum/EPP model.^26^ In that paper the authors assumed that under their accelerated scale-up ART coverage only reaches 85% of those with a CD4^+^ cell count below 350/uL or about 50% of all those infected with HIV and that ART only reduces infectivity of HIV patients by 80%. If we make similar assumptions our model is able to reproduce the results of the Spectrum/EPP model for Mozambique. This shows that there is still a need to reach consensus among modellers on the levels of coverage, compliance and viral load suppression that can be reached.

## Appendix 3. Cost of the death of a young adult

Estimating the cost to the state of the death of a young adult is difficult and there does not appear to be a consensus on how best to do this. For example it has been argued that ‘total annual treatment costs, including drugs and medical services, are now around $500 to $1,000 per year in sub-Saharan Africa, probably about the same as the average annual income of the prime-aged workers being struck down by the disease… making [this] cost effective’.^33^ The implication is that the cost of treatment should be less than the average earnings. This in turn would suggest that the cost to society of the death of a young adults is equal to the life-time average earnings. The average *per capita* gross wage in South Africa is US$14k^34^ roughly double the GDP *per capita*^35^ so that the cost of a death would be about US$420k. If this were the case then the cost of deaths would exceed all other costs by orders of magnitude.

One might rather want to consider only the revenue lost to the national fiscus. If the government receives about 10% of wages in the form of taxes the loss of revenue when a person dies will be about US$1.4k *p.a*. but with unemployment running at about 25% they can presumably be replaced within a year or so and the cost of a death would only be US$1.4k or about 20% of GDP.

Another approach is to consider the cost of in-patient hospitalization for those that then die. In the countries of East and Southern Africa each day spent in a primary level hospital costs 0.8% (Range: 0.66%–1.1%) of GDP *p.a*. and 1.0% of GDP *p.a*. in South Africa.^36^ In Baragwanath Hospital in South Africa the mean time from admission to death for those that died of AIDS in the hospital was 10 days^37^ giving a cost of hospitalization of US$577 per death or 8% of GDP *p.a*.

If we are only concerned with the costs to the state of the death of a working age man or woman then it would seem that a cost of 15% of GDP *p.a*. may be reasonable. If we were to include the costs to the state beyond the health services, mainly from loss of wages, the need to retrain and recruit new workers, care for orphaned children, as well as other costs arising from the disruption of the social fabric, and so on, the cost could be significantly higher.

## Appendix 4: Comparison with projections made by Walensky *et al*.^25^

In a related study Walensky^25^ has estimated the impact of treatment on the HIV epidemic and the costs of treatment under a *current pace strategy*, roughly equivalent to our, *constant effort strategy*, and the *UNAIDS strategy*, roughly equivalent to our *expanded treatment and prevention strategy*. In the Walensky analysis^25^ the number of people living with HIV in South Africa remains unchanged over the next ten years at about 6.6 million under the *current pace strategy* and drops to about 6 million under the *UNAIDS strategy*. Our analysis yields similar results with number of people living with HIV over the next ten years increasing from 6.4 million to 7.0 million under the *constant effort strategy* and remaining steady at 6.3 million under the *expanded treatment and prevention strategy*, allowing for population growth. However, the two studies differ in important respects. Over the next ten years there will be 4.4M new infections and 5.0M deaths under the Walensky *current pace strategy*^25^ but only 1.9M new infections and 0.8M deaths under our *constant effort strategy*. Over the next ten years there will be 2.4M new infections and 2.5M deaths under the Walensky *UNAIDS strategy* but only 0.4M new infections and 0.2M deaths under our *expanded treatment and prevention strategy*.

In 2016 about 6.3M adults were living with HIV in South Africa of whom about 2.7M are not on ART. Without treatment about 10% of them will die each year. If there is no further increase in treatment coverage; then there should be of the order of 2.7M new infections and about the same number of deaths over the next ten years. The Walensky estimate for the current pace strategy would seem to be too high by a factor of about 2 and their estimates for the UNAIDS strategy would be about right if expanded treatment had no impact on new infections or deaths. Resolving these differences should be a priority.

## Appendix 5. Cost comparison with Walenski *et al*.^25^

Walensky *et al*.^25^ use costs which are not dissimilar to those in Table 3. They have US$137 and US$375 *p.a*. for first and second line treatment. This would give the same estimate as used here if 42% were on first and 58% on second line ART. They have US$7 and US$20 per test for HIV-negative and HIV-positive people, respectively, about half of our estimate in Table 3 of US$32. (Médicine sans Frontières provide a recent study of costs estimates for a range of tests.^38^) Walensky *et al*.^25^ use costs of US$ 20, 27, 32, 70, 157 for providing health care to people in clinical stages 1 to 4 of diseases and with terminal illness, respectively, which are similar to the values used in Table 3. They also use a cost of US$ 770 for tuberculosis, similar to ours, and include costs of treating other bacterial and fungal diseases. They include US$155 *p.a*. for adherence and retention on ART which is not included in our analysis.

The total costs of the *current pace* and *constant effort strategies* over the next ten yeas are estimated to be about US$38Bn and US$54Bn, respectively. Our model suggests that the total costs of the *constant effort* and *expanded treatment and prevention strategies* over the next ten years are estimated to be US$21 Bn and US$25 Bn, respectively. The Walensky estimates of the number of incident cases, deaths and total costs are roughly double ours so that this discrepancy needs to be resolved. It should be noted that our model gives a total cost for managing HIV and TB in 2013 of 2013 US$2.2Bn which is close to the value of ZAR22 Bn given in the SANAC Investment Case report.^17^

